# CXCL12 chemokine dimer signaling modulates acute myelogenous leukemia cell migration through altered receptor internalization

**DOI:** 10.1101/2024.08.26.609725

**Authors:** Donovan Drouillard, Michael Halyko, Elizabeth Cinquegrani, Maria Poimenidou, Miracle Emosivbe, Donna McAllister, Francis C. Peterson, Adriano Marchese, Michael B. Dwinell

## Abstract

Acute myeloid leukemia (AML) is a malignancy of immature myeloid blast cells with stem-like and chemoresistant cells being retained in the bone marrow through CXCL12-CXCR4 signaling. Current CXCR4 inhibitors that mobilize AML cells into the bloodstream have failed to improve patient survival, likely reflecting persistent chemokine receptor localization on target cells. Here we characterize the signaling properties of CXCL12-locked dimer (CXCL12-LD), a bioengineered variant of the naturally occurring oligomer of CXCL12. CXCL12-LD, in contrast to wild-type or locked monomer variants, was unable to induce chemotaxis in AML cells. CXCL12-LD binding to CXCR4 decreased G protein, β-arrestin, and intracellular calcium mobilization signaling pathways, and indicated the locked dimer is a partial agonist of CXCR4. Despite these partial agonist properties, CXCL12-LD increased CXCR4 internalization compared to wildtype and monomeric CXCL12. Analysis of a previously published AML transcriptomic data showed CXCR4 positive AML cells co-express genes involved in survival, proliferation, and maintenance of a blast-like state. The CXCL12-LD partial agonist effectively mobilized stem cells into the bloodstream in mice suggesting a potential role for their use in targeting CXCR4. Together, our results suggest that enhanced internalization by CXCL12-LD partial agonist can avoid pharmacodynamic tolerance and may identify new avenues to better target G protein coupled receptors.

## INTRODUCTION

The chemokine CXCL12 can self-associate into dimers at high concentrations or following interactions with its cognate receptors or glycosaminoglycan binding partners [1]. To determine the impact of ligand oligomerization on chemokine function the Volkman group has engineered CXCL12 into variants locked into either a dimeric structure, CXCL12-locked dimer (CXCL12-LD) or a monomer form, CXCL12-locked monomer (CXCL12-LM) [2]. These forms are structurally identical to native CXCL12 conformations attained by oligomerization of the wild-type protein and retain the ability to bind to and activate the cognate receptor CXCR4 or the atypical chemokine receptor 3 (ACKR3) [3]. Recognizing that chemotaxis typically follows a biphasic response CXCL12-LD and CXCL12-LM were used to uncover discrete functional roles for the monomer in stimulating cell migration, compared to the dimer, which induced a stationary phenotype we termed “ataxis” [4, 5]. Thus, our data from cell culture models suggest that CXCL12 induced chemotaxis, with maximal migration observed in a narrow concentration range of ligand, reflects formation of dimers at high concentrations. *In vivo*, CXCL12-mediated chemotaxis regulates the trafficking of CXCR4-expressing hematologic cells into primary and secondary lymphoid tissues, angiogenesis, and is a key component in solid cancers for promoting metastasis [6, 7]. In hematological malignancies, CXCR4 is thought to function less in chemotactic migration and more as a retention signal that keeps malignant cells within the bone marrow where they are protected from chemotherapy [8]. While we have previously dissected CXCL12 signaling in colon, breast, and pancreas solid tumors [9–11], the pharmacologic properties of monomeric and dimeric ligands in hematological malignancies, such as acute myeloid leukemia (AML), remains unknown.

AML is a heterogenous malignancy of adults and children characterized by clonal expansion of immature myeloid blast cells with resulting bone marrow failure and ineffective erythropoiesis [12]. The standard of care chemotherapy regimen during induction therapy consists of daunorubicin and cytarabine. This regimen is relatively effective in younger patients with a 5-year survival rate of 55% [13], but ineffective in older patients with a 5-year survival rate of only 17% in patients greater than 60-years-old due to a lower tolerance for high-dose chemotherapies [14]. With a median diagnosis age of 68 years [15], there is a pressing need to develop new, non-cytotoxic therapies to treat AML in the elderly. An attractive therapeutic target in the leukemia microenvironment is the CXCL12-CXCR4 chemokine axis, which helps maintain AML cells in their protective bone marrow niche [16]. AML cell lines variably increase CXCR4 expression upon chemotherapy treatment, resulting in increased CXCL12-mediated chemotaxis and conferring a bone marrow stroma-mediated survival advantage [17]. In preclinical models with human AML xenografts, AMD3100, a clinically available inhibitor of CXCR4 known by the trade name Plerixafor, combined with chemotherapy demonstrated chemosensitizing effects supporting a role for the CXCL12-CXCR4 axis as a mediator of stromal-dependent chemo-resistance [18, 19]. In addition, a number of small peptide CXCR4 antagonists have been studied in preclinical mouse models and have shown similar chemosensitizing effects through a variety of pathways including mobilization of AML cells into the peripheral blood [20], activation of pro-apoptotic pathways [21, 22], or inhibition of proliferation [20]. Despite this preclinical success, phase I/II clinical trials combining AMD3100 with a variety of chemotherapy regimens in adults with AML have found little to no clinical benefit [22–25].

Clinical trials with the newer generation of CXCR4 inhibitors were, as with AMD3100, terminated due to their lack of efficacy [26, 27]. The failure of current CXCR4 inhibitors highlights the need to better understand the signaling mechanisms whereby CXCL12 and CXCR4 regulate AML functions. Using rationally designed and engineered CXCL12 variants locked into their dimeric structure we have uncovered that the dimer is a partial agonist of CXCR4 that stimulates ligand-induced receptor internalization that is stronger and more sustained compared to monomer or wild-type ligand. This internalization occurs independent of chemotaxis as the dimer was unable to stimulate cell migration, despite retention of G protein signaling and β-arrestin recruitment. The sustained internalization of CXCR4 by dimeric ligand suggests a new pathway to disrupt CXCL12-mediated retention of AML within the protective bone marrow niche.

## RESULTS

### Human AML cell lines are CXCR4+ and migrate to CXCL12-WT but not CXCL12-LD

As a first step to investigate mechanisms of chemokine signaling in AML cells, we used RT-PCR and flow cytometry to assess the expression of CXCR4, the cognate receptor for CXCL12 in three human AML cell lines. Consistent with a prior report, CXCR4 mRNA (**Fig. 1A**) and protein (**Fig. 1B**) was highly expressed in three different AML cells [28]. Transcript for ACKR3, an atypical chemokine receptor CXCL12 may also bind, was also detected on each of the AML cell lines (**Supplemental Fig. 1A**).

**Figure 1:**
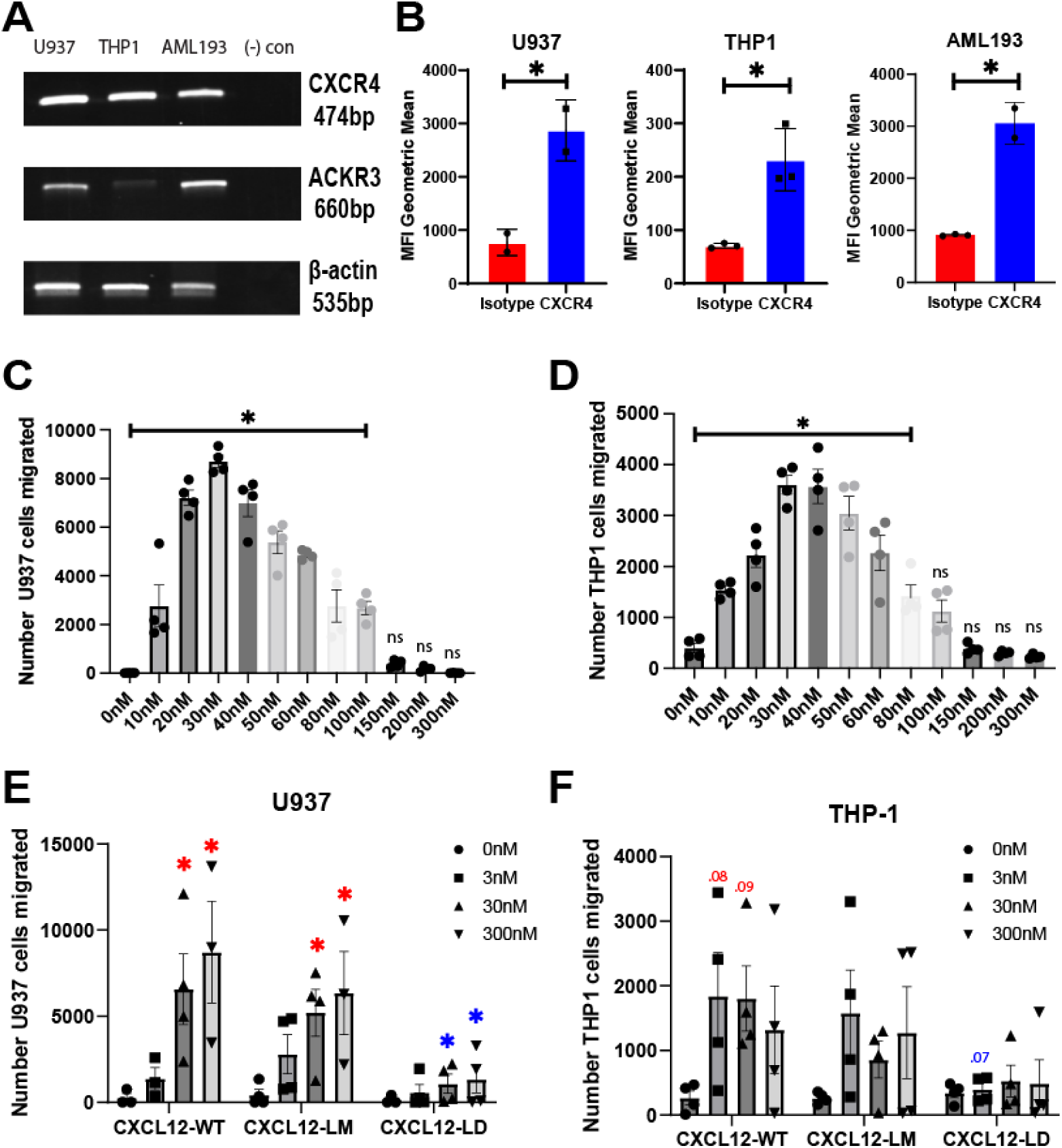
CXCR4 expression in AML cell lines. **(A)** RT-PCR of CXCR4, ACKR3, and beta actin in three human AML cell lines U937, THP-1, and AML-193. Data representative of 3 separate analyses. **(B)** Flow cytometry results of the three AML cell lines stained with either anti-CXCR4 clone 12G5 or control isotype antibody. * = *P* ≤ 0.05 by Student’s t-test. n = 2-3 independent analyses. Values are mean ± SEM. **(C)** U937 or **(D)** THP-1 cell chemotaxis from the top chamber of a transwell filter towards the lower chamber containing the indicated concentration of CXCL12-WT chemokine. All values under significance bar represent a significant increase by one-way ANOVA with Dunnett’s multiple comparison test to 0 nM vehicle treated. **(E)** Total number of U937 or **(F)** THP-1 migrated from the top chamber of a transwell chamber towards the lower chamber containing the indicated concentration of CXCL12-WT, CXCL12-LM, or CXCL12-LD. Migrating cells in panels C-F were enumerated after a two hour incubation using flow cytometry. Values represent mean ± SEM, n = 3-4 independent biological replicates. Red asterisks or numbers indicate adjusted p value by two-way ANOVA with Tukey’s multiple comparisons test to that respective variant’s 0 nM (*e.g.* CXCL12-WT 30 nM to CXCL12-WT 0 nM) while blue indicates significance compared to the same dose of a different variant (*e.g.* CXCL12-LD 30nM to CXCL12-WT 30nM).

Accumulating evidence from our lab and others support a model wherein the biphasic chemotactic migration induced by chemokines may reflect distinct binding modes of monomeric and dimeric forms of the chemokine. In the CXCL12 model, chemotaxis reflects receptor activation by ligand monomers, with dimer formation becoming predominant as the concentration of the ligand increases, shifting the cells to ataxis. Consistent with that model, AML cells undergo dose dependent migration in response to CXCL12-WT protein with an optimal concentration of approximately 30 nM (**Fig. 1C**, **1D**). Concentrations above 80 nM resulted in a less robust chemotactic response, consistent with the expected bell-shaped chemotactic curve due to possible CXCL12-WT dimerization at higher concentrations. In contrast to wild-type or monomeric variants, data from transwell migration assays in two separate AML cell lines indicate that CXCL12-LD was unable to stimulate migration at 3, 30, or 300 nM concentrations (**Fig. 1E**, **1F**). However, we did not observe a difference between CXCL12-WT and a variant locked into a monomer and unable to form dimers, CXCL12-LM [10].

Prior CXCL12 structure-function analysis established a substantial role for the first two amino acids in the ligand amino terminus, Lys1 and Pro2, in receptor activation, with a complete loss of Ca^2+^ flux agonist activity upon deletion or substitution of either residue [29]. Previous work with a CXCL12 variant lacking these first two residues (CXCL12_3–68_) suggest that this molecule binds to CXCR4, albeit with reduced affinity [11, 29]. Consistent with those data, CXCL12_3-68_ failed to induce chemotaxis at any concentration (**Supplemental Fig. 1B, 1C**).

### CXCL12 G protein signaling and arrestin recruitment

CXCR4 activates heterotrimeric G proteins, leading to inhibition of adenylyl cyclase and mobilization of intracellular calcium. As shown in **Figure 2**, we first assessed G_αi_ signaling in THP-1 cells treated with CXCL12-WT, CXCL12-LD, or CXCL12_3-68_. Consistent with its binding to G_αi_-coupled CXCR4, CXCL12-WT inhibited cAMP production in a dose-dependent manner with significant inhibition occurring at a concentration of 1000 nM (**Fig. 2A**). While the dimer was unable to promote chemotaxis, CXCL12-LD inhibited cAMP production at 1000 nM **(Fig. 2A**, dark grey bar) while CXCL12_3-68_ failed to activate the receptor (**Fig. 2A, light grey bar**).

**Figure 2:**
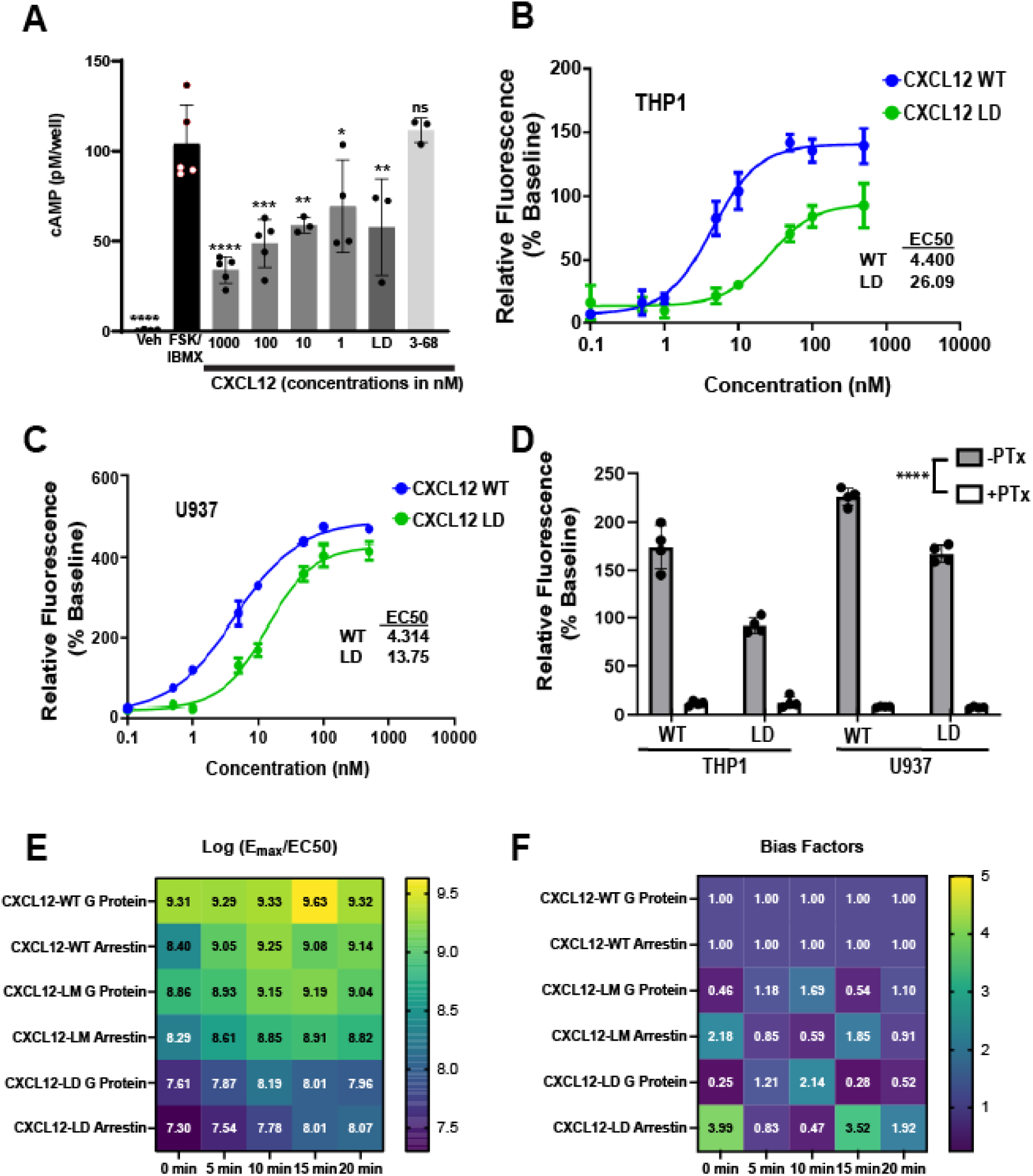
CXCL12-LD is a balanced partial agonist. **(A)** cAMP levels in THP-1 cells stimulated with 10 mM IBMX and 1 μM forskolin followed by treatment with various compounds measured by a competitive enzyme immunoassay. All significance values are in relation to FSK/IBMX by two-way ANOVA with Dunnett’s multiple comparison test. **(B, C)** Calcium flux in THP-1 and U937 cells treated with either CXCL12-WT or CXCL12-LD. n = 3. **(D)** AML cells were pre-treated with pertussis toxin prior to treatment with CXCL12 variants. All pertussis toxin pre-treated values are significantly different by Student’s t-test when compared to the same form of CXCL12 without pertussis toxin. **(E)** The log(E_max_/EC_50_) of the CXCL12 variants from net BRET in HEK293-CXCR4-Luc cells transfected with β-arrestin or Gαi Venus transducer. n = 6, EC_50_ values calculated from non-linear curve fit, all r^2^ values > 0.82. **(F)** Bias factors for CXCL12-WT, -LM, and -LD at all timepoints tested from HEK293 BRET studies calculated using the calculator found on the Biased Signaling Atlas [30]. A bias factor >5 is indicative of biased agonism.

To analyze another output of GPCR activation we measured intracellular calcium mobilization. We found that both CXCL12-LD and CXCL12-WT induced intracellular calcium flux in THP-1 and U937 AML cell lines (**Fig. 2B**, **2C**). CXCL12-LD induced calcium flux but with less potency and efficacy than CXCL12-WT. To confirm if calcium flux was mediated by activation of the G_αi_ subunit of the GPCR, we performed the same experiment on cells treated in the presence or absence of pertussis toxin, which ADP-ribosylates G_αi_ to prevent the activation of the G protein. CXCL12-induced calcium flux was extinguished by the addition of pertussis toxin (**Fig. 2D**), demonstrating ligand activation of CXCR4 mediates calcium flux through activation of the G_i/o_ family of proteins.

In addition to G protein signaling, CXCL12 binding to CXCR4 induces β-arrestin recruitment. Using bioluminescence assays, we measured the recruitment of β-arrestin to CXCR4 following treatment with either CXCL12-WT, CXCL12-LM, or CXCL12-LD. Compared with wild-type or monomer ligand, CXCL12-LD recruited β-arrestin with less potency over a 20 minute time course (**Supplemental Fig. 2, left panels**). Similarly, using a mini-G_αi_ BRET we confirmed the lower calcium mobilization evoked by CXCL12-LD and found that the dimer less potently activated the CXCR4 G protein complex (**Supplemental Fig. 2, right panels**). Data from each of the individual G-protein and arrestin BRET experiments were then used to calculate the transduction efficiencies for each (**Fig. 2E**). Lastly, the bias factors were calculated for each ligand, with a bias factor of ≥5 indicative of a biased agonist [30]. The calculated bias factors demonstrated that CXCL12-LD signals as a balanced partial agonist while CXCL12-LM is a balanced full agonist in relation to CXCL12-WT (**Fig. 2F**) in AML cells.

β-arrestin recruitment requires phosphorylation of Ser/Thr residues in the CXCR4 C-terminus. The intracellular CXCR4 C-terminus has 18 Ser/Thr residues whose site-specific phosphorylation is dynamically regulated by G protein regulatory kinases (GRKs) [31]. As a first step, we used a phospho-specific antibody against dually phosphorylated Ser324 and Ser325 (Ser324/325) of the CXCR4 C-terminal tail. Compared to vehicle treated CXCR4 transfectants (**Fig. 3A**), cells treated with 10 nM CXCL12-WT (**Fig. 3B**) or CXCL12-LM (**Fig. 3C)** had increased CXCR4 phosphorylation at Ser324/325. In contrast, CXCL12-LD resulted in weak CXCR4 C-terminal serine phosphorylation (**Fig. 3D**). Altogether, these data support the notion that CXCL12-LD is a partial agonist that has lower potency and efficacy at signaling through either G_αi_ and β-arrestin.

**Figure 3:**
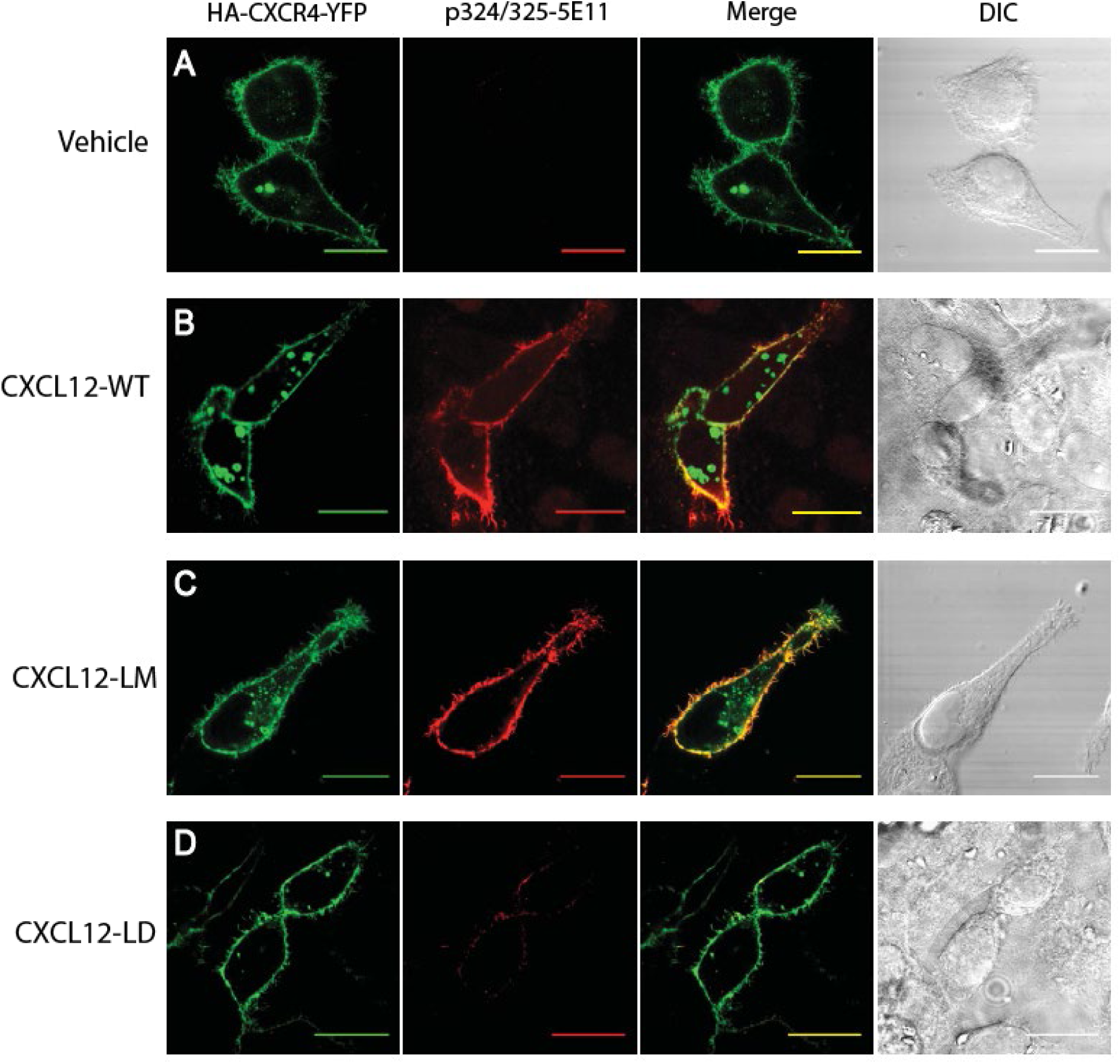
Differential CXCR4 C-terminal phosphorylation in response to CXCL12 variants. **(A)** HeLa cells transiently transfected with HA-CXCR4-YFP were stimulated with vehicle, **(B)** CXCL12-WT, **(C)** CXCL12-LM, or **(D)** CXCL12-LD. The far left panels show visualization of YFP-CXCR4 and the middle left panel is visualization of the phosphorylated Ser324/325 residues on CXCR4. Co-localization of the YFP-CXCR4 and phosphorylated CXCR4 appear yellow in the merge panel on the middle right. Differential image contrast (DIC) images are shown on the far right panels. Shown are the representative micrographs from three independent experiments. Bars = 20 μm.

### CXCL12-LD internalizes CXCR4 more effectively than CXCL12-WT

Canonical GPCR internalization occurs largely through β-arrestin dependent signaling mechanisms and has been measured for CXCL12-WT in a variety of cells including human T cells and the rat basophilic leukemic cell line RBL-2H3 [32, 33]. Current antagonists of CXCR4 signaling do not internalize the receptor, likely reflecting their functioning as competitive inhibitors occupying a key ligand binding site [33–35]. Despite the dimer’s limited recruitment of β-arrestin, we measured a significant dose-dependent decrease in CXCR4 surface localization in cells treated with CXCL12-LD at both 30 minutes and 24 hours (**Fig. 4A**, **4B**). The 12G5 monoclonal antibody binds CXCR4 at the 2^nd^ extracellular loop, a process that may be blocked when different agonists are bound to the receptor [36]. To more stringently evaluate if the receptor was being internalized, we repeated the experiment using monoclonal antibody clone 1D9, an antibody that binds the N-terminus of the receptor and can therefore detect both ligand-bound and un-bound CXCR4 [37]. Just as with 12G5, we measured decreased surface levels of CXCR4 after 30 minutes (**Fig. 4C**) and 24 hours (**Fig. 4D**) of CXCL12-WT, CXCL12-LM, or CXCL12-LD treatment. Both 12G5 and 1D9 antibodies detected a statistically significant decrease in CXCR4 on the surface of cells treated for 24 hours with CXCL12-LD, but not CXCL12-LM, compared to CXCL12-WT treated cells. Moreover,1D9 measured a significant internalization of CXCR4 at 30 minutes that was sustained through 24 hours. CXCR4 internalization was next measured in THP-1 and AML193 cell lines treated with 100 nM CXCL12-WT, CXCL12-LM, or CXCL12-LD and confirmed the larger and more sustained internalization induced by the dimer variant (**Fig. 4E**, **4F**). Lastly, as we had shown decreased CXCR4 C-terminal phosphorylation of S324/325 needed for arrestin-receptor interactions, we measured CXCR4 internalization in CXCR4-FLAG transfected HeLa cells. Consistent with our data in AML cells, antibody against the N-terminal FLAG tag confirmed increased internalization with CXCL12-LD relative to wild-type protein or CXCL12-LM (**Supplemental Fig. 3**).

**Figure 4:**
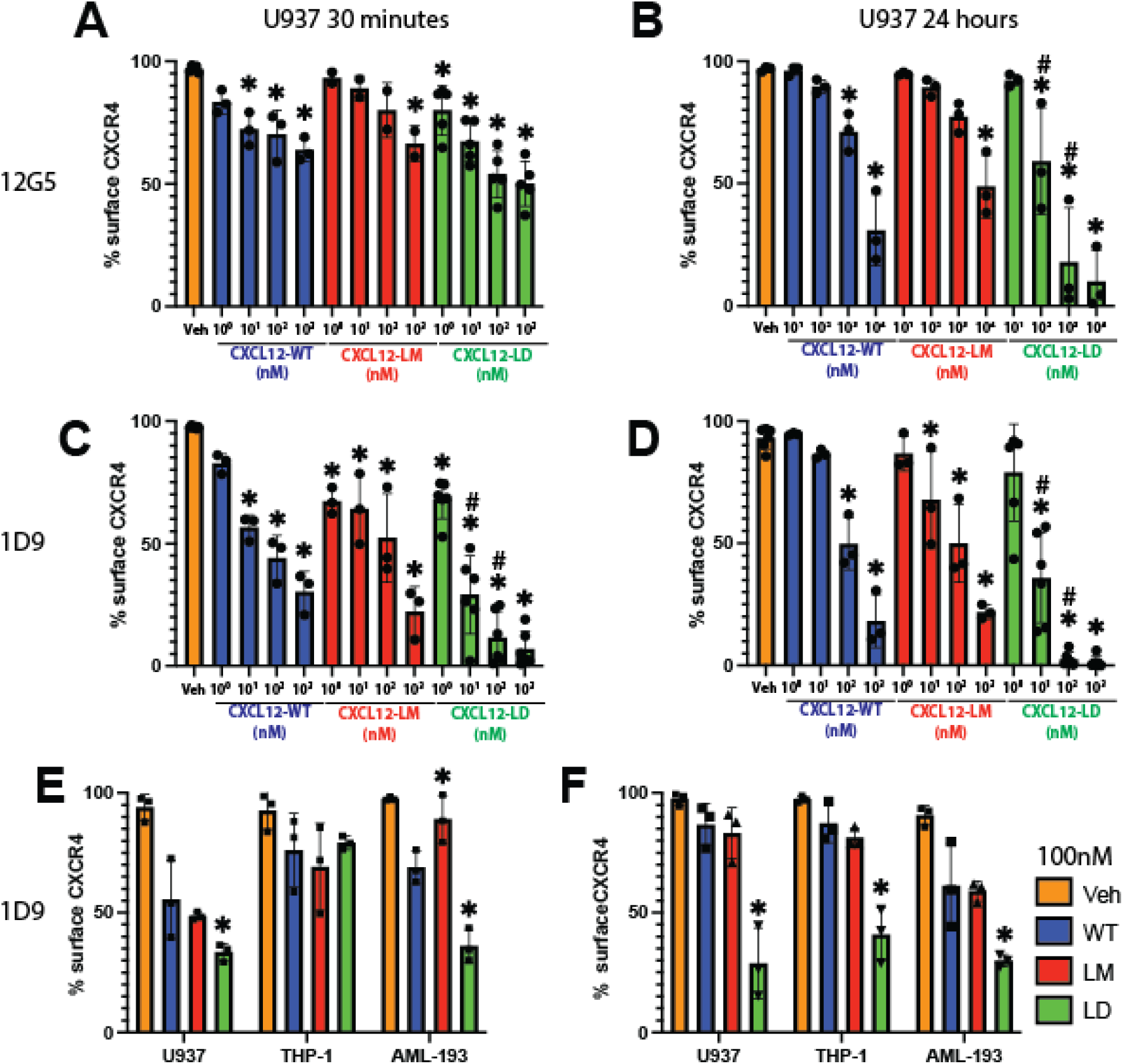
CXCL12-LD promotes internalization of CXCR4. **(A)** Percent of CXCR4 positive U937 cells detected by flow cytometry with the 12G5 CXCR4 antibody after 30 minutes or **(B)** 24 hours of incubation with various concentrations of CXCL12 WT (blue), LM (orange), or LD (green). * represent significance when comparing to vehicle and # represents significance when comparing to respective dose of CXCL12-WT (*e.g.* CXCL12-LD 10 nM to CXCL12-WT), adjusted *P* ≤ 0.05 by two-way ANOVA with Tukey’s multiple comparison test. **(C)** The percent of CXCR4 positive U937 cells detected by flow cytometry with the 1D9 CXCR4 antibody after 30 minutes or **(D)** 24 hours of incubation with various concentrations of CXCL12 variants. **(E)** Three different leukimic cell lines treated with CXCL12 WT, LM, or LD and stained with the 1D9 antibody after 30 minutes of treatment or **(F)** 24 hours. Significance represents comparison of CXCL12-variant to CXCL12-WT in the same cell line by one-way ANOVA with Dunnett’s multiple comparison test, *P* ≤ 0.05.

As the three AML cell lines express both CXCR4 and ACKR3, we next asked if the ligand variants modified internalization of the atypical chemokine receptor. ACKR3 functions as a scavenger receptor that helps sculpt the chemokine gradient without directly invoking cellular migration [38] and is an arrestin-biased receptor in response to CXCL12 [39]. Unlike the internalization measured for CXCR4, none of the CXCL12 oligomers induced internalization of ACKR3 in U937, THP-1, or AML-193 cells (**Supplemental Fig. 4**).

As an additional measure to verify that the decreased surface CXCR4 staining measured by flow cytometry was not due to steric hindrance of the binding site by CXCL12-LD, we generated CXCL12 variants with AzDye647 conjugated to the ligands C-terminal tail. U937 cells treated with fluorescent dye tagged CXCL12-WT, CXCL12-LM, or CXCL12-LD for 30 minutes demonstrated robust internalization of the ligand (**Fig. 5A-C**). Significantly more CXCL12-LD-AzDye647 was detected within U937 cells compared to wild-type or monomeric chemokine (**Fig. 5D**). These fluorescence microscopy data support the notion that CXCL12-LD promotes receptor internalization rather than sterically hindering CXCR4 antibodies from binding.

**Figure 5:**
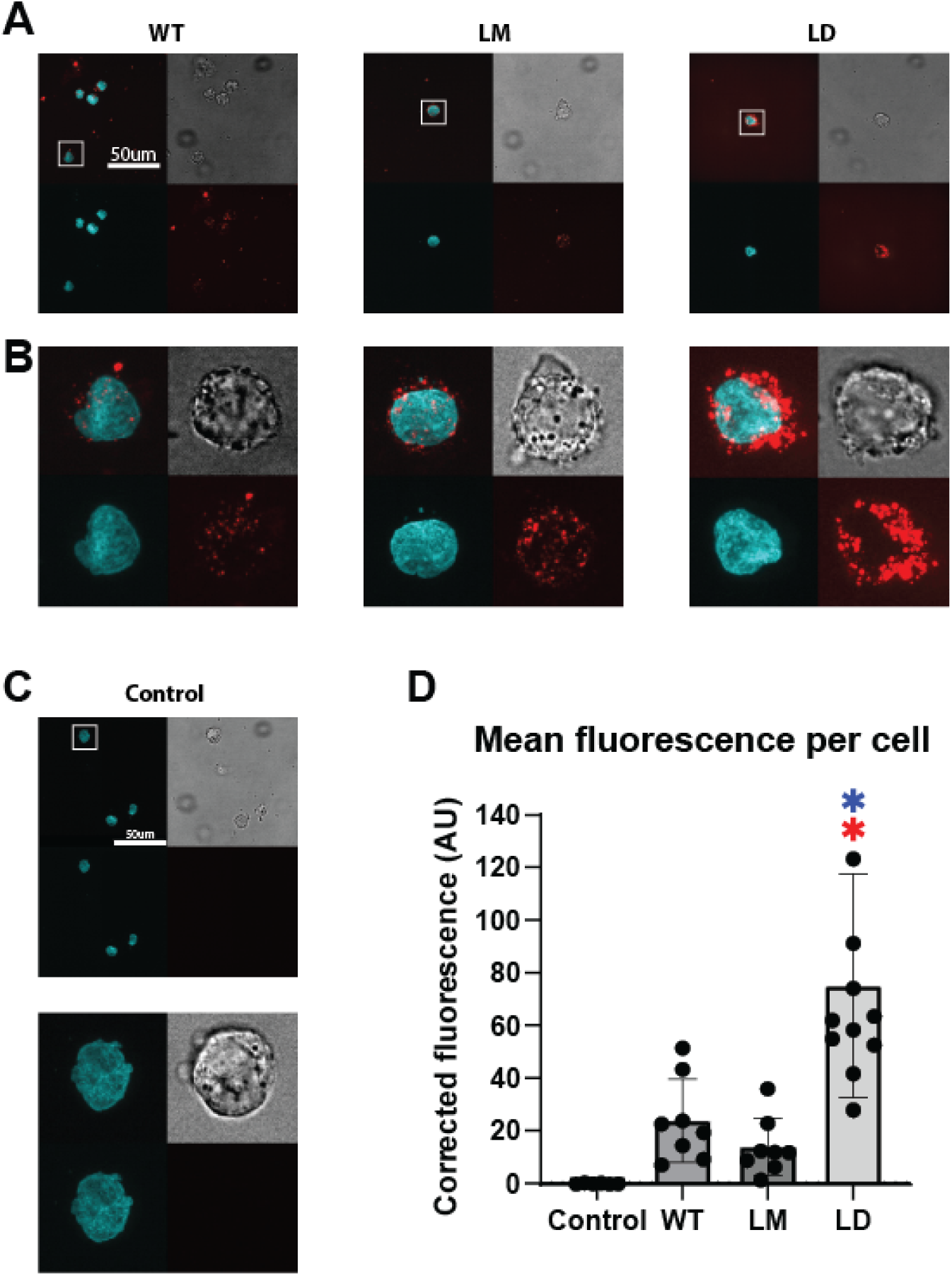
Labeled CXCL12 ligands show internalization with CXCL12-LD. **(A)** U937 cells were treated with 200 nM of AzDye647-conjugated CXCL12 variants (red, bottom right of one image) of their respective variant for 30 minutes before fixation and staining with Hoechst 3342 (cyan, bottom left). Brightfield is shown in the top right and the merged image is the top left. Images shown are at 100X. Scale bar is 50 μm. **(B)** Enhancement on the cutout of the images in the white square from Fig. 5A. **(C)** Untreated U937 cells shown under the same conditions as figure 5A and 5B. **(D)** Quantification of the corrected (background substracted) average red AzDye647 fluorescence intensity inside of cells. Red asterisks indicate significance compared to control and blue asterisks indicate signficance compared to CXCL12-WT by one-way ANOVA with Tukey’s multiple comparisons test, *P* ≤ 0.05.

### Partial agonist CXCL12-LD but not CXCR4 antagonists stimulate receptor internalization

Previous reports suggest CXCR4 small molecule antagonists induce therapeutic resistance when used to treat patients [23–27]. Conventional wisdom is that therapeutic resistance reflects an upregulation of the receptor by the antagonist [40–44]. To test that notion, CXCR4 surface levels were measured in U937 (**Fig 6A**, **6B**) or AML-193 (**Fig 6C**, **6D**) cells treated with the receptor antagonists AMD3100 or BL-8040 and compared against cells incubated with CXCL12-LD. CXCR4 levels were increased in cells treated with AMD3100 or BL-8040 at both the 30 minute and 24 hour treatment (**Fig. 6A**, **6B**). AML-193 cell showed a more modest increases in surface CXCR4 after 24 hours of treatment with AMD3100 or BL-8040 (**Fig. 6C**, **6D**). These data are in stark contrast to CXCL12-LD treated cells, in which CXCR4 surface levels were significantly reduced in both U937 and AML-193 cells at both time points.

**Figure 6:**
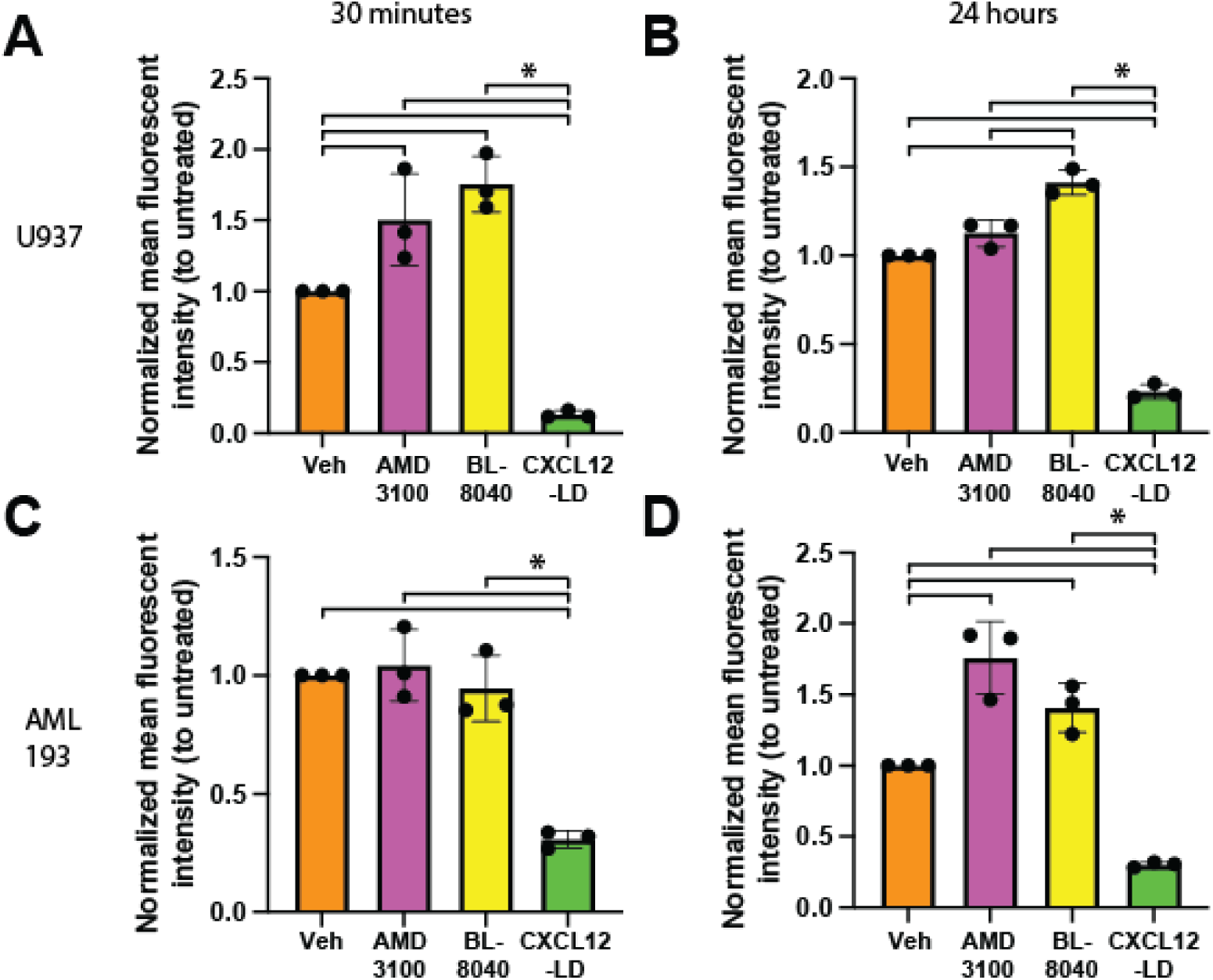
CXCR4 surface upregulation seen with AMD3100 and BL-8040 but not CXCL12-LD. **(A)** U937 cells were treated with either 1 μM AMD3100, 100 nM BL-8040, or 100 nM CXCL12-LD and immunostained for surface CXCR4 with the 1D9 clone. Mean fluorescent intensity (MFI) values were normalized against vehicle-treated (PBS) U937 cells stained for surface CXCR4. Treatment was for 30 minutes or **(B)** 24 hours. **(C)** CXCR4 inhibitor treatment of AML-293 cells for 30 minutes or **(D)** 24 hours under the same conditions seen in 6A and 6B. * = *P* ≤ 0.05 by one-way ANOVA with Tukey’s multiple comparisons test.

### CXCR4 is upregulated in chemoresistant AML blasts

The unique ataxic and internalizing properties of CXCL12-LD prompted further investigation into the potential for the dimer to influence AML *in vivo*. CXCR4 is important for trafficking and retention of hematopoietic cells in the bone marrow [45]. To analyze the CXCR4 expressing cells in AML patients, we turned to publicly available large single-cell RNA datasets. Using a dataset that contains bone marrow aspirates of 16 AML patients [46], we found CXCR4 expression was enriched myeloid lineage cells (**Fig. 7A**). To examine if CXCR4 was upregulated in malignant monocytes compared to healthy monocytes, we compared the monocyte-like cells, which have a confirmed mutation in targeted sequences known to be mutated in AML, to healthy, non-mutated monocytes from the same donors. CXCR4 expression was found to be upregulated in the malignant monocytes compared to the healthy monocytes (**Fig. 7B**). Next, we investigated transcriptional differences between CXCR4 positive and negative malignant monocytes in AML patients. CXCR4+ cells had upregulated expression of the transcription factor JUN and transcriptional coactivator CITED2, each of which are necessary for AML blast survival [47–49] (**Fig. 7C**). Further, HLX and SAP30, two proteins important for blocking differentiation and enhancing proliferation, respectively, were similarly elevated in malignant CXCR4+ AML [50–52]. These gene correlations were confirmed with the AML PanCancer Atlas dataset on cBioPortal (**Supplemental Fig. 5**). Lastly, as some transcription factors are lowly expressed and thus difficult to detect by single cell RNA analysis [53], we completed analyses using the CollecTRI R package [54]. ETV5 and FOXH1 were within the top results in the CXCR4+ group, both of which are important for preventing differentiation and maintaining a blast-like state in AML [55, 56] (**Fig. 7D**). Together these data suggest that CXCR4 is upregulated in AML and that CXCR4+ AML blasts similarly upregulated genes associated with resisting apoptosis, proliferation, and maintaining and a stem-like state.

**Figure 7:**
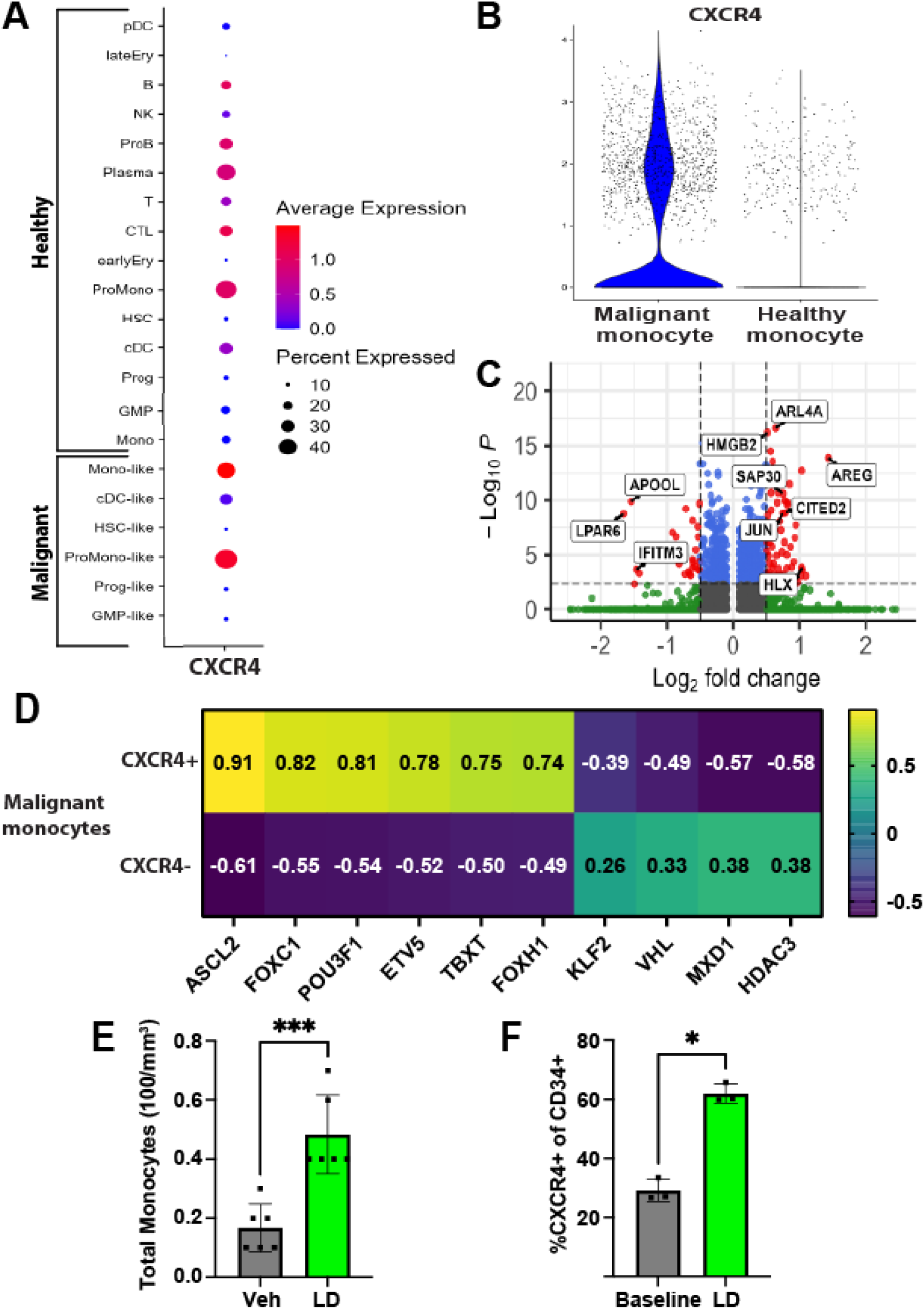
CXCL12-LD increases the frequency of hematopoietic progenitor cells in peripheral blood. **(A)** Dot plot of CXCR4 expression in the bone marrow aspirates of 16 AML patient’s pre-treatment [46]. Each of the “-like” cells had a confirmed mutation in a targeted DNA sequence to confirm their malignancy. **(B)** The “Mono” and “Mono-like” cell types were further analyzed for CXCR4 expression. CXCR4 was elevated in the Mono-like (Malignant monocyte) compared to the Mono (Healthy monocyte) by MAST test (*P* ≤ 0.05) [84]. **(C)** The malignant monocyte population from figure 6B was grouped by CXCR4 expression. Differentially expressed genes by MAST test were found and plotted on the volcano plot. Grey dots represent non-significant genes by both log2FC or adjusted p-value, green dots represent genes with significant adjusted p-value but not log2FC, blue dots represent genes with significant log2FC but not adjusted p-value, and red dots represent genes with both significant log2FC and adjusted p-value. Log2FC threshold was 0.5 and adjusted p-value threshold was 0.005. **(D)** The malignant monocytes sorted by CXCR4 expression were analyzed using the CollecTRI R package to estimate the transcription factor activity based on weighted mean gene expression with the obtained t-values of the slope plotted. Positive values represent inferred transcription factor activity while negative values represent inferred inactive transcription factor activity. **(E)** Wildtype C57BL/6 mice were given 200 μL 5 μM CXCL12-LD or vehicle PBS. The frequency of total monocytes **(E)** and CD34+ CXCR4+ progenitor cells **(F)** in peripheral blood was measured two hours after treatment. Values are mean ± SD from 3-6 separate mice.

### CXCL12-LD mobilizes monocytes and CXCR4+ hematopoietic stem cells in mice

Small molecular receptor antagonists of CXCR4 are hypothesized to chemosensitize AML cells through mobilization of AML blasts into the peripheral blood [18, 26, 49]. AMD3100 was the first bicyclam CXCR4 antagonist shown to increase the concentration of circulating hematopoietic stem cells in mouse models [19]. To date, however, AMD3100 has had limited success in clinical trials of AML, resulting in no improvement in remission rates compared to historical controls and indicating a need for further exploration of CXCR4 therapeutic targeting [26]. To determine if chemokine oligomers can mobilize CXCR4-expressing cells *in vivo*, wild-type C57BL/6 mice were injected subcutaneously with CXCL12-LD and blood was collected after two hours to assess monocyte mobilization and after four hours to assess CXCR4+ hematopoietic stem cell mobilization. We observed an increase in circulating monocytes in CXCL12-LD treated mice compared to vehicle treated mice (**Fig. 7E**). Further, there was an increase in the percentage of CXCR4+ circulating hematopoietic stem cells (**Fig. 7F**). These data support the hypothesis that CXCL12-LD promotes long-term mobilization of CXCR4-expressing cells into the peripheral blood and could be used to remove AML blasts out of the chemoprotective bone marrow niche.

## DISCUSSION

Cumulatively, our data support the summary model in **Figure 8** suggesting that retention of hematopoietic or transformed cancer cells in the bone marrow may be disrupted by the unique signaling properties of the partial agonist CXCL12-LD. Using reductionist AML cell culture models, we determined that CXCL12-LD, compared to wild-type or monomeric ligand, is a partial agonist of CXCR4. We observed increased internalization of CXCR4 and not ACKR3 with CXCL12-LD and a complete inability to evoke chemotaxis. In comparison to other inhibitors of CXCR4 chemotaxis, AMD3100 is a 1^st^ generation CXCR4 inhibitor that likely functions as a competitive antagonist and both LY2510924 and BL-8040 (Motixafortide, T140, BKT140) are 2^nd^ generation CXCR4 inhibitors that function as inverse agonists with higher potency compared to AMD3100 [57, 58]. While AMD3100, BL-8040, and CXCL12-LD all block chemotaxis, only CXCL12-LD reduced surface CXCR4 expression while both AMD3100 and BL-8040 increase it. As both chemotaxis and receptor internalization have been canonically linked with β-arrestin signaling [32] our data suggest a new mechanism for homologous CXCR4 desensitization and a potential avenue for avoiding resistance to CXCR4 inhibitors.

**Figure 8:**
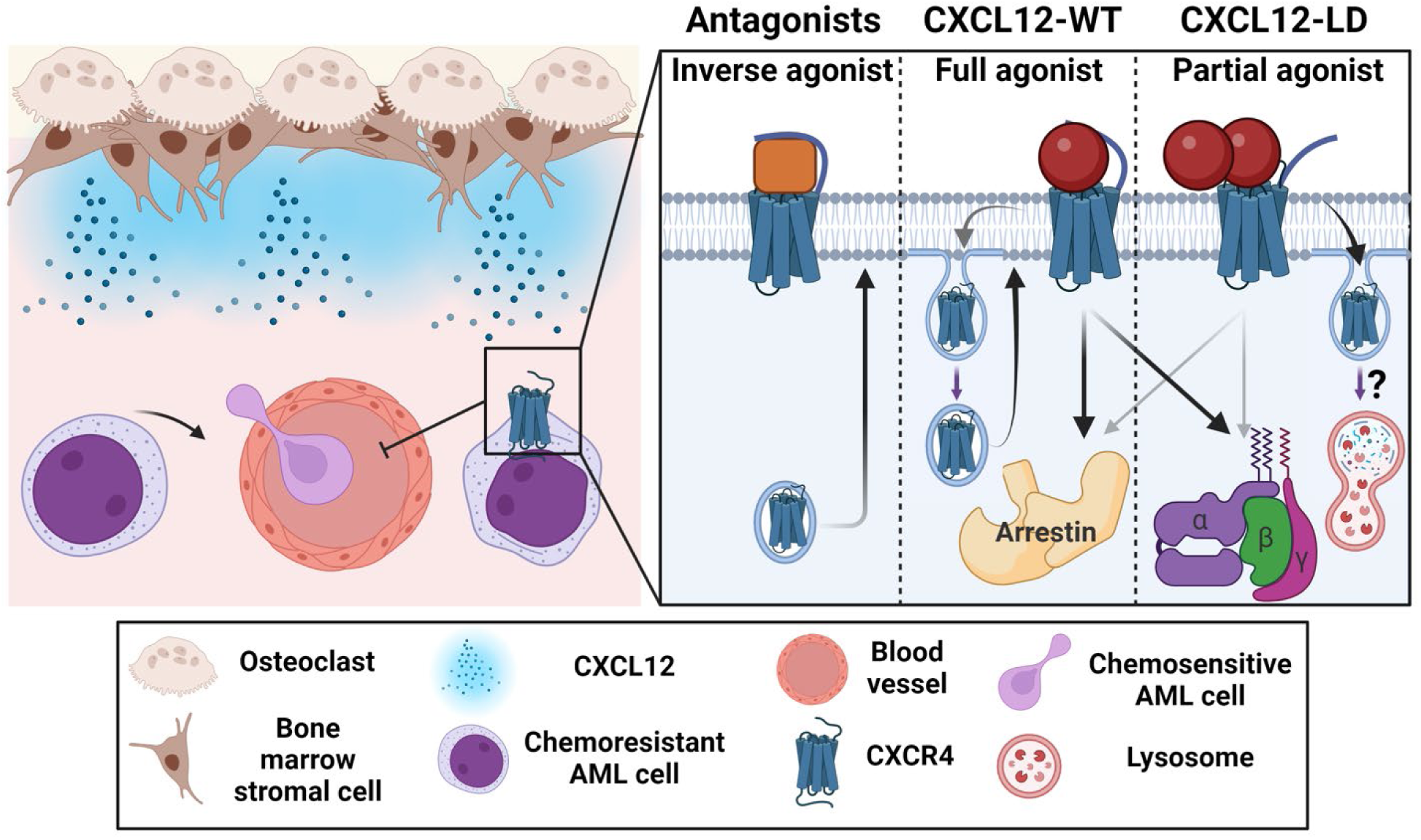
CXCL12-LD functions differently than currently available CXCR4 inhibitors. **Left**, CXCL12 produced by bone marrow stromal cells maintains chemoresistant CXCR4-expressing AML blasts in the marrow. If CXCR4 expression is lost or the receptor is inhibited, the blasts are mobilized into the bloodstream. **Right**, 2^nd^ generation CXCR4 inhibitors are shown to be inverse agonists of CXCR4 that inhibit internalization and result in surface receptor upregulation. In the middle, CXCL12-WT as a balanced agonist activating both β-arrestin and the G-protein signaling pathways. Signaling with CXCL12-WT results in receptor internalization and recycling. On the right, CXCL12-LD is a partial agonist whose signaling results in greater receptor internalization compared to CXCL12-WT, possibly through increased receptor lysosomal degradation rather than endosomal recycling. Created in BioRender. Drouillard, D. (2025) https://BioRender.com/t28xc88.

Prior studies of CXCL12-WT show it engages with CXCR4 at four different sites on the receptor, or chemokine recognition sites (CRS) [59]. CXCL12-WT binds CRS0.5, 1.0, 1.5, and 2.0 while CXCL12-LD can bind all but CRS0.5 [60, 61]. However, the inability to bind CRS0.5 is thought to only affect signaling efficacy, suggesting that CRS0.5 is required for full ligand agonism [60]. The lack of CXCL12-LD binding to CRS0.5 supports the idea that the decreased CXCR4 1D9 antibody binding measured is not due to steric hindrance by the CXCL12-LD agonist, as the 1D9 antibody clone binds the flexible N-terminus and CRS0.5 is within the distal N-terminus of the receptor [62]. Another distinction between CXCL12-LD and CXCL12-WT binding occurs when examining post-translational CXCR4 tyrosine sulfation. Both CXCL12-LD and CXCL12-WT can bind sulfated residues Tyr-21 and Tyr-12 on the CXCR4 amino terminus, while CXCL12-LD differs from wild-type ligand by binding to sulfated Tyr-7 [3, 63]. Additionally, sulfated Tyr-21 binds CXCL12-LD with 20-fold stronger affinity than CXCL12-WT or CXCL12-LM and sulfated Tyr-21 promotes ligand dimerization [3]. These discrete differences in ligand oligomer interactions with CXCR4 may help explain the discrepancy between cellular migration and receptor internalization. CXCL12-LD binds the receptor in a dimerized state, while CXCL12-WT likely binds as a monomer and then dimerizes [3]. This difference in sequential binding and dimerization may explain why dimer binding to CXCR4 stimulates pronounced and durable internalization in the absence of chemotactic migration.

Another quandary posed by our data is the increased internalization with CXCL12-LD despite decreased β-arrestin recruitment. β-arrestin independent internalization of GPCRs has been observed previously and can occur through multiple mechanisms *via* clathrin coated pits [64]. New studies suggest that CXCR4, but not all GPCRs, are internalized through sorting nexins (SNX) and particularly SNX9 and SNX18 [65]. This process was shown to be partially driven by the phosphorylation of residues on the proximal CXCR4 C-terminus. The phosphorylation sites required to recruit SNX9 are different from those required for β-arrestin, in line with our data suggesting reduced β-arrestin recruitment but increased internalization by CXCL12-LD. While we show decreased phosphorylation of residues Ser324/325, we did not examine Ser338/339 in the proximal cluster which could be driving SNX9 recruitment and CXCR4 internalization. Further, β-arrestin binding and SNX9 binding to CXCR4 are thought to be mutually exclusive [65]. A decrease in β-arrestin binding, as seen with CXCL12-LD, could allow for increased SNX9 binding and internalization.

Overexpression of CXCR4 has been demonstrated in a variety of hematologic and solid cancers including AML [17, 66, 67]. While preclinical experiments implicate CXCR4 antagonism as a promising means of chemosensitizing AML and other hematologic malignancies, clinical results have shown limited efficacy [27, 67, 68]. A shared feature of these CXCR4 antagonists is blockade of ligand binding and downstream functional activity. As shown here, AMD3100 and other small molecule antagonists known to inhibit receptor signaling may paradoxically increase receptor surface levels, which in turn negates the benefit sought in blocking receptor signaling, a process known as “antagonist tolerance”. Antagonist tolerance can also be seen in other GPCRs and may be a key contributor to treatment resistance and the lack of clinical benefit seen with AMD3100 [40–44, 69–71]. Extrapolating from those *in vitro* data would suggest that AML tolerance to CXCR4 inhibition through persistent surface localization of receptor occurs within a crucial time-window preceding the completion of the initial week-long induction chemotherapy. Using an engineered protein to dissect pharmacological signaling of CXCR4 we have extended our initial discovery of partial agonism in pancreas cancer cells to a high-risk hematologic malignancy to examine how agonists of the same receptor function. The differential ability of CXCL12-LD to promote receptor internalization provides a potential avenue to address roles in pharmacotherapy that traditional antagonists have been unable to address.

## MATERIALS and METHODS

### Cell Lines

Human U937 (CRL-1593.2) and THP-1 (TIB-202) cells were obtained from the ATCC (Manassas, VA) and maintained in RPMI with L-glutamine and 25 mM HEPES (Life Technologies Inc, Carlsbad, CA) supplemented with 10% (v/v) fetal bovine serum (Omega Scientific, Tarzana, CA). HEK-293 epithelial cells (CRL-1573) and HeLa cells (CCL-2) purchased from the ATCC were maintained in DMEM (Life Technologies Inc) supplemented with 10% (v/v) fetal bovine serum (Omega Scientific). Human AML-193 (CRL-9589) cells were obtained from the ATCC and maintained in IMDM supplemented with 5% (v/v) fetal bovine serum, 1X Insulin-Transferrin-Selenium (Gibco), and 5 ng/mL GM-CSF (R&D Systems). Cell lines were authenticated annually using short tandem repeat profiling and mycoplasma-tested semi-annually. HEK293 cells were transfected with HA-CXCR4 using TransIT-LT1 transfection reagents (Mirus Bio, Madison, WI) as previously described [72]. HeLa cells were transiently transfected using polyethylenimine as previously defined [73].

### Recombinant Chemokines

CXCL12 wild-type (CXCL12-WT), locked monomer (CXCL12-LM), locked dimer (CXCL12-LD) and NH_2_-terminal truncation (CXCL12_3-68_) variants were expressed and purified as previously described [10, 74]. The CXCL12-LD mutations are L36C and A65C and create a dimeric protein [2, 11]. The CXCL12-LM mutations are L55C and I58C [10, 69]. The engineered locked dimer or locked monomer proteins are structurally indistinguishable from native dimeric and monomeric protein, respectively, and retain full CXCR4 receptor binding capability [4, 10]. Proteins were expressed in *E. coli* and bacterial cells lysed by French press. Fusion protein was purified through nickel chromatography, refolded by infinite dilution, and ULP1 protease was used to cleave the 6XHIS-Sumo tag. CXCL12-WT and CXCL12-LD were fluorescently labeled with AzDye647 using a two-step process employing sortagging [75] and copper-catalyzed azide-alkyne click chemistry as defined previously [76]. The identity of native or fluorescently labeled CXCL12 was confirmed by linear ion trap quadrupole mass spectrometry. Lyophilized proteins were stored at −20°C.

### Flow Cytometry

THP-1, U937, and AML-193 cells were passaged to 5 x 10^5^ cells/mL the day prior to experiment. Cells were washed twice using cold PBS, incubated in Fc blocking solution (Miltenyi Biotec, Bergisch Gladbach, Germany), and washed twice. Cells were then incubated for 30 minutes in fluorophore conjugated primary CXCR4 antibody (Invitrogen, Waltham, MA) on ice and washed twice. Fluorescence intensity was measured on a BD-LSR II flow cytometer and analyzed by FlowJo software (BD Biosciences, San Jose, CA).

### RT-PCR

RNA was isolated using the RNA Easy kit from Qiagen and treated with DNAse I to remove genomic DNA contaminants. Conversion to cDNA was performed by priming with random hexamers and the SuperScript II cDNA synthesis kit (Life Technologies). PCR products were separated by agarose gel electrophoresis and visualized by ethidium bromide staining and ultraviolet imaging. Primers for actin were used as a loading control. Amplification of chemokine receptors was done using the following primers (5′-3′):

Human CXCR4 Forward 5’-GACTCCATGAAGGAACCCTGTTTCCG-3’

Human CXCR4 Reverse 5’-CTCACTGACGTTGGCAAAGATGAAGTCG-3’

Human β-actin Forward 5’-ACCCACACTGTGCCCATCTACG-3’

Human β-actin Reverse 5’-AGTACTTGCGCTCAGGAGGAGC-3’.

Human ACKR3 Forward 5’-GTGGTGGTCTGGGTGAATATC-3’

Human ACKR3 Reverse 5’-GATGTAGTGCAGGATGGAGAAG-3’

### cAMP Quantification

THP-1 cells were grown in a T-75 flask, then counted immediately prior to the measurement of cAMP using a competitive enzyme immunoassay as we have shown before [77]. THP-1 (5 x 10^5^) cells were aliquoted into 1.5 mL tubes and stimulated with 10 mM 3-isobutyl-1-methylxanthine (IBMX) for 30 minutes at room temperature, after which they were incubated with either CXCL12-WT, CXCL12-LD, or CXCL12-3-68 for an additional 10 minutes. Cells were then stimulated for 10 minutes with 10 μM forskolin (FSK) to stimulate cAMP production. Controls included unstimulated cells and cells treated with IBMX and FSK alone. Cells were then centrifuged and incubated with 0.1 M HCl for 20 minutes before being centrifuged (300 x g on microcentrifuge) for 10 minutes. Supernatant was isolated, and cAMP was assayed using a competitive enzyme immunoassay kit (Cayman Chemical Co., Ann Arbor, MI) according to the manufacturer’s instruction manual. Fluorescence was measured at room temperature with a SpectraMax Multimode Microplate reader (Molecular Devices, San Jose, CA) at a wavelength of 450 nm. Readings were graphed and analyzed according to the immunoassay instruction manual.

### Calcium Mobilization

Calcium mobilization was measured using the FLIPR Calcium 6 assay kit (Molecular Devices) according to manufacturer’s directions. Cells were plated to 50,000 cells/well in 96-well plates. Cells were washed with Calcium/Magnesium-free HBSS supplemented with 20 mM HEPES (Thermo Fisher Scientific, Waltham, MA) and 0.1% (v/v) BSA Fraction V (Thermo Fisher Scientific) and loaded with Calcium-6 dye for 1 hour at 37°C and 5% CO_2_. Chemokines were diluted in calcium/magnesium-free HBSS buffered with 20 mM HEPES and were then loaded onto a separate compound plate. Fluorescence was measured at 37°C with a FlexStation3 Multimode Microplate Reader (Molecular Devices) with excitation and emission wavelengths of 485 and 515 nm, respectively. Chemokines were resuspended at the concentrations indicated in the figure legends and added to the cells after baseline fluorescence was measured for 20 seconds. The percentage Ca^2+^ flux was calculated from the maximum fluorescence minus the minimum fluorescence as a percent of baseline fluorescence. EC_50_ values were determined by nonlinear fitting to a four-parameter logistic function. For experiments with addition of the Gα subunit inhibitor, cells were treated with 100 ng/mL pertussis toxin for 4 hours prior to assessing calcium flux in response to 500 nM CXCL12-WT or CXCL12-LD.

### Confocal Immunofluorescence Microscopy

Phosphorylated CXCR4 serine residues 324 and 325 were detected in HeLa cells as described previously [78], grown on 10 cm dishes were transfected with 5 μg of HA-CXCR4-YFP and passed 24 hours later onto coverslips pretreated with poly-l-lysine, as described previously [79]. Cells were washed once with DMEM plus 20 mm HEPES and serum-starved for 3-4 hours in the same medium, followed by 30-minute treatment with vehicle PBS, 80ng/mL CXCL12-WT, CXCL12-LM, or CXCL12-LD at 37°C. Following treatment, cells were fixed, permeabilized, and immunostained using a custom mouse monoclonal antibody specific for dually phosphorylated serine residues 324 and 325 (pSer324/325) (Clone 5E11), as described previously [78, 80]. Images were acquired using a Zeiss LSM 510 laser-scanning confocal microscope with a 63x W Apochromat oil-immersion objective. Image acquisition settings across parallel samples were identical.

### Transwell Migration

Cells were passaged to 5 x 10^5^ cells/mL the day prior to individual experiments. On the day of the experiment, cells were counted, washed twice, and plated to the upper well of a Transwell insert (Corning Incorporated, Corning, NY) at a density of 60,000 cells/well in assay medium (RPMI-1640, 0.2% (w/v) BSA Fraction V (Thermo Fisher Scientific), with chemoattractants added to the bottom well in assay medium. Cell migration was measured after 2 hours. Transwell migration was enumerated by flow cytometry using the High Throughput Sampler attachment for the BD LSR-II flow cytometer. Results were analyzed with FlowJo software (BD Biosciences).

### BRET Assays

HEK293T cells were passaged to 2 x 10^6^ cells/mL the day prior to the experiment. On the day of the transfection, cell media was changed one hour prior to the introduction of the transfection mixture. OptiMEM was warmed to 37°C and 500 mL was aliquoted into a microfuge tube, along with 0.06 μg Receptor-CXCR4-RLuc8. 5 μg Venus-Transducer β-arrestin and MiniG, and 15.9 μL TransIT. The transfection mixture was incubated in the cell culture hood for 20 minutes and then added dropwise to the cell plates. Cells were then placed back into the incubator overnight.

The day of the BRET assay, cells were washed with PBS and trypsinized for 5 minutes. DMEM supplement with 10% (v/v) FBS was then added to neutralize the trypsin before cells were centrifuged for 5 minutes at 300 x g. Cells were then resuspended in PBS + 0.1% (v/v) glucose. A 15 μL aliquot of cells was placed into a microfuge tube and counted using hemocytometer. Cells should be at a final density of 100,000 cells/well of a 96-well plate. Cells were then plated into a 96-well plate and incubated for 1 hour. Next, the ligand plate was prepared, with the ligands of interest being CXCL12-WT, -LM, and –LD. Ligands were diluted in PBS + 0.1% (v/v) glucose to reach the desired final concentration of 30 μM. Serial dilutions of the ligands were then made in the 96-well plate. Coelenterazine H was resuspended to 1 mM using methanol, and a 50 μL working stock of Coelenterazine H into PBS + 0.1% (v/v) glucose buffer was made. 100 μL of this working stock was added to each well in row A of the 96-well plate. 10 μL of the Coelenterazine H was then transferred from row A to the rest of the rows of the 96-well plate. The Omega plate reader (BMG Labtech; Cary, NC) was then used to measure bioluminescence. Net BRET was calculated using Excel.

### Receptor Internalization

U937, THP-1, and AML-193 cells were counted and placed into 5mL polystyrene tubes at 10^6^ cells/mL in 1.0 mL. Cells were treated with the denoted concentrations and variants of CXCL12 for the indicated times. After, the cells were washed twice with 0.5% (v/v) BSA in PBS. The cells then underwent a Fc receptor block (Biolegend, 422302) for 10 minutes, washed twice with 0.5% (v/v) BSA in PBS buffer, and then cell surface stained with either PE-conjugated CXCR4 antibody clone 12G5 (Invitrogen, 12-9999-42) or PE-conjugated clone 1D9 (BD, 551510) for 30 minutes. ACKR3 surface levels were measured using ACKR3-APC (Biolegend, 331114). After staining, the cells were washed twice with flow buffer and resuspended in 300-500 μL flow buffer. Surface CXCR4 and ACKR3 expression was quantified by analyzing the stained cell suspension on a Cytek Aurora spectral flow cytometer. AMD3100 was used at a concentration of 1 μM and BL-8040 was used at 100 nM. Results were analyzed with FlowJo software.

### CXCL12-AzDye647 Imaging

2.5 x 10^5^ U937 cells were grown on 24-well glass-bottom plates pre-coated with poly-D-lysine to enhance attachment. The cells were then treated with one of the following conditions: Vehicle-Control, 200 nM of CXCL12-WT-AzDye647, CXCL12-LM-AzDye647, or CXCL12-LD-AzDye647 for 30 minutes. After treatment, cells were counterstained with Hoechst nucleic acid stain to visualize nuclei and fixed with 4% (w/v) paraformaldehyde for 15 minutes. Imaging was performed using the Andor BC43 microscope (Oxford Instruments, Abingdon, UK) at 100X. Images were analyzed using ImageJ and Amaris software. To quantify average fluorescence intensity per cell, the brightfield images of the cells were used to trace the cell border of all cells within the image. An area containing no cells was used to subtract background fluorescence from (**Supplemental Fig. 6**). The ImageJ EzColocalization plugin [81] was used to calculate the average red fluorescence intensity. The red channel only image was set as channel one with the cell-traced regions of interest set as the cell identifier. If multiple cells were within one image, the average fluorescence intensity per cell within that image was averaged before subtracting the background fluorescent intensity. The quantification shown represent an N of 2-3 with multiple 100X images taken per well with each dot representing the results from one image.

### Bioinformatic Analysis

Analysis of a previously published dataset containing 16 AML patients and five healthy donors [45] was performed using the Seurat v5 (5.1.0) package in R version 4.4.1 [82]. The patient data was integrated using the Harmony package [83] Transcriptional regulon analysis was performed using the CollecTRI R package [53]. The volcano plot was generated using the EnhacedVolcano package. Single-cell RNA analysis of differentially expressed genes was performed using the MAST test [84]. The code for the analysis can be found at https://github.com/The-Michael-Dwinell-Lab-at-MCW/CXCL12-Locked-Dimer-AML. cBioPortal was used for gene correlations from The Cancer Genome Atlas. Specifically, the acute myeloid leukemia dataset was selected from the Pan-Cancer dataset with CXCR4 as the gene query.

### Mice and Bone Marrow Cell Mobilization

All experiments were conducted under approved protocols from the Medical College of Wisconsin (AUA00076) and in accordance with the National Institutes of Health Guide for the Care and Use of Laboratory Animals and the ARRIVE guidelines [85]. Wildtype C57BL/6J were obtained from The Jackson Laboratory (Bar Harbor, ME). Animals were maintained on a strict 12:12 hour light–dark cycle in a temperature- and humidity-controlled facility with water and food provided ad libitum. Mice were injected with 200 μL 5 μM CXCL12-LD or 200 μL PBS subcutaneously in the right flank. Two hours after injection, mice were euthanized, and blood collected by cardiac puncture. Isolated blood was analyzed on the scil-Vet-abc Hematology Analyzer (scil Animal Care Company GmbH, Viernheim, Germany) to calculate the total number of monocytes per mm^3^. In separate groups of mice, blood was collected from submandibular bleed 4 hours after injection with 200 μL 5 μM CXCL12-LD subcutaneously in the right flank. Red blood cells were lysed using manufacturers protocol (Biolegend 420301) and hematopoietic cells stained with Fixable viability dye eFluor 455 (BD Biosciences 65-0868-14), murine Fc block (Bio X Cell BE0307), murine anti-CD45 PerCP (ThermoFisher MA1-10234), murine anti-CXCR4 SuperBright 600 (ThermoFisher 63-9991-80), and/or murine anti-CD34 FITC (ThermoFisher 11-0341-82) and enumerated on a BD Fortessa flow cytometer (BD Biosciences, Franklin Lakes, NJ).

### Ligand Transduction Coefficients and Bias Calculations

Raw values obtained from individual BRET experiments for both the G protein and β-arrestin recruitment were used to calculate a transduction ratio and bias relative to CXCL12-WT at each time point tested as previously described [84]. To calculate transduction coefficients, CXCL12-LM and CXLC12-LD values were normalized to CXCL12-LD. The Emax and EC_50_ values were calculated using the obtained normalized values. The values were then used to calculate the transduction ratio, or log of the relative activity using log RA = log(Emax/EC_50_). The control-normalized factor was calculate using Δlog RA = log RA _CXCL12 isoform_ – log RA _CXCL12-WT_. The log of the bias factor was calculated using ΔΔlog RA _CXCL12 isoform_= log RA_Pathway1_ – log RA_Pathway2_. Lastly, the final bias factor was calculated using 10^ΔΔlog^ ^RA^ _CXCL12 isoform_ [30].

### Interalization in HeLa cells by ELISA

HeLa cells grown in 6-cm dishes were transiently transfected with CXCR4 (1 μg) tagged with FLAG and N-luc on the N- and C-terminus, respectively. The next day, 2 x 10^5^ cells were seeded into a 24-well plate pre-coated for at least 1 hour with 0.1 mg/mL poly-L-lysine (Sigma; cat. No. P1399). Following overnight incubation at 37°C, cells were rinsed once with 500 μl of serum-free DMEM and treated with or without ligand for 30 minutes at 37°C. Cells were rinsed once with 500 μL ice-cold TBS buffer (20mM Tris, pH 7.5, 150mM NaCl) and immediately fixed for 5 minutes with 500 μL of 3.7% (v/v) formaldehyde diluted in TBS (Sigma; cat. No. F8775). Cells were washed three times with 500 μl TBS followed by incubation for 45 minutes at room temperature in TBS supplemented with 0.1% (w/v) bovine serum albumin (BSA) (Gold Bio cat. No. A-420-10). Thereafter, cells were incubated for 1 hour at room temperature with anti-FLAG M2 alkaline phosphatase-conjugated antibody diluted 1:3000 in TBS-BSA. Cells were washed 3 times with TBS and then incubated with a developing solution for 20 minutes when a light-yellow color was formed. Aliquots of each reaction were transferred to a 96-well plate containing 0.4 M of NaOH to quench the reaction. Absorbance readings were measured at 405 nm using a microplate reader. After background subtraction, receptor internalization was determined as a loss of surface receptor after ligand stimulation relative to the vehicle.

### Statistical Analyses

Statistical analyses were performed using Prism 8.0 software (GraphPad Software, Irvine, CA). Power analysis was performed using an alpha error probability of 0.05 and a power level of 0.8 to select rigorous sample sizes for individual experiments. Unpaired sample comparisons between two groups were analyzed by Student’s t-test when data was normally distributed with equal variances of the groups, or by Mann-Whitney test when parametric test conditions were violated. Three or more independent groups were compared using one-way or two-way ANOVA with Dunnett’s or Tukey’s HSD test for multiple comparisons (test used denoted in figure legends).

## ACKNOWLEDGMENTS

We thank Galina Petrova, PhD for expert guidance and training in the Children’s Research Institute Flow Cytometry Shared Resource. We gratefully thank Megan C. Harwig, PhD, Microscopy Core Manager, for her invaluable assistance in completing the immunofluorescence microscopy.

## Author contributions

Author Contributions (CRediT Roles):

Conceptualization: MH, EC, DD, MBD

Data curation: MH, DM, DD

Formal Analysis: DD, MH, EC, DD

Funding acquisition: MBD

Investigation: DD, MH, EC, DM, AM, MP, ME

Methodology: DD, MP, MH, AM, MBD,

Project Administration: MBD

Resources: AM, FP, MBD

Software: AM

Supervision: MBD

Validation: MH, DM

Visualization: DD, MH, MP

Writing – original draft: MH

Writing – review and editing: DD, MH, EC, FP, AM, MBD

## DATA AVAILABILITY

This study did not generate new unique reagents. All data are contained within the manuscript.

