## Supplemental Data for "CXCL12 chemokine dimer signaling modulates acute myelogenous leukemia cell migration through altered receptor internalization"

**Supplemental Figure 1**

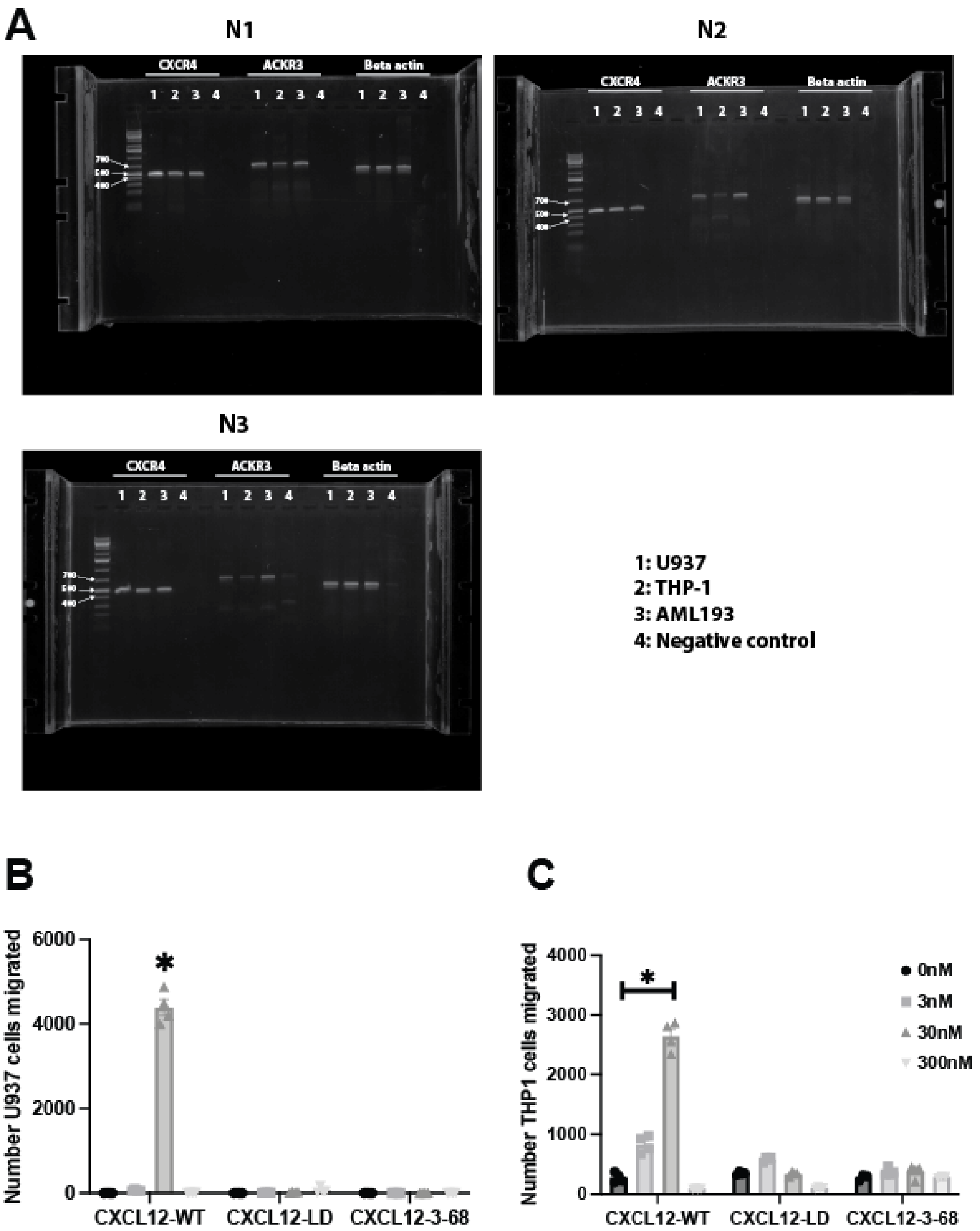


**AML cell lines express CXCR4 and ACKR3. (A)** Three biological replicates of PCR testing for CXCR4 (left), ACKR3 (middle), with beta actin as a positive control (right). The band order is U937, THP-1, AML193, and a negative control. The gels shown are unedited. **(B)** Total number of U937 or **(C)** THP-1 migrated from the top chamber of a transwell chamber towards the lower chamber containing the indicated concentration of CXCL12-WT, CXCL12-LD, or CXCL12-3-68. Migrating cells in were enumerated after a two hour incubation using flow cytometry. All significance values are in relation to the ligands 0nM control cells by one-way ANOVA with Dunnett’s multiple comparison test.

**Supplemental Figure 2**

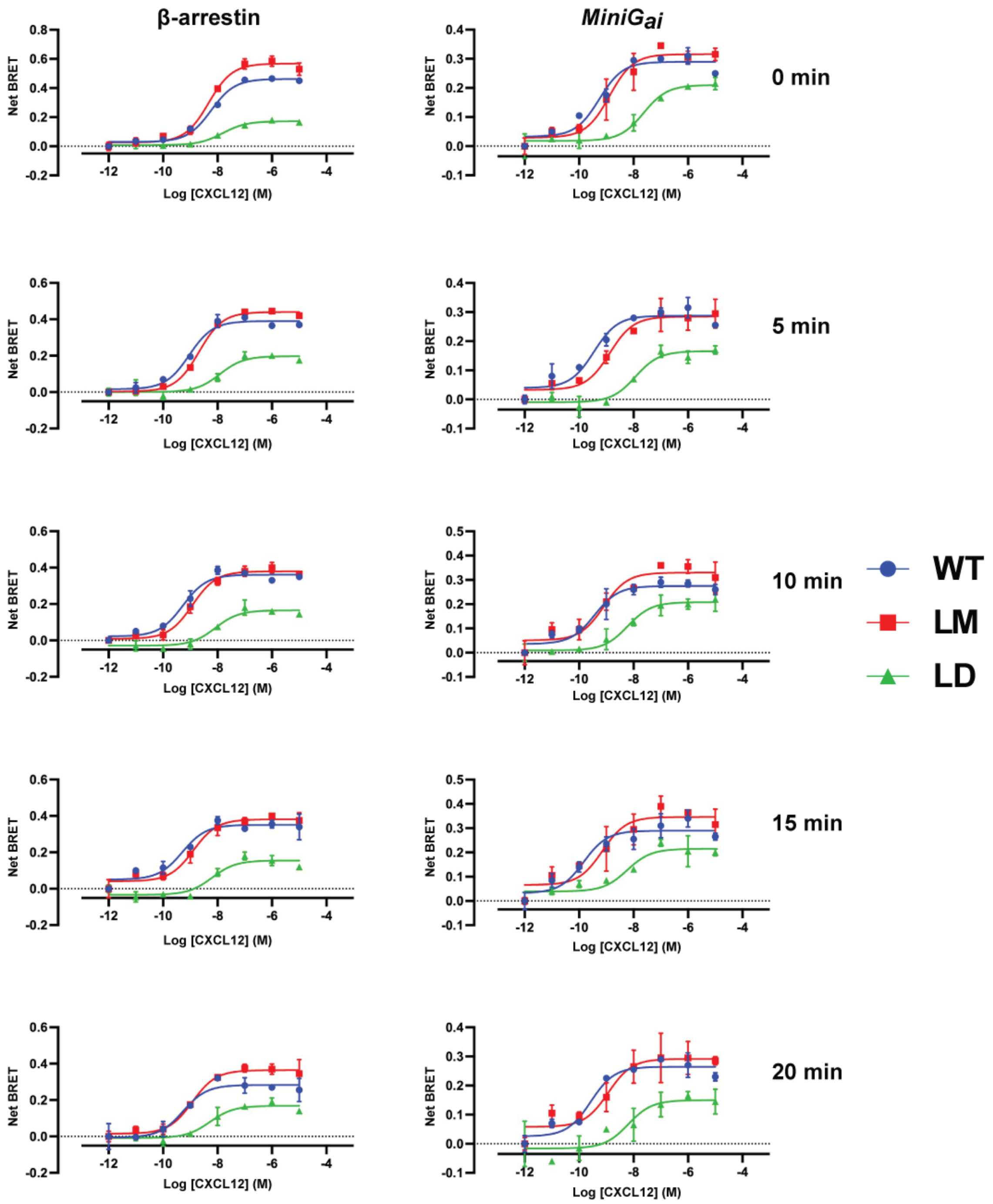


**G protein and β-arrestin recruitment with CXCL12 variants.** Net BRET in HEK293-CXCR4-Luc cells transfected with β-arrestin or Gαi Venus transducer. N = 6. Time points indicate length of incubation with the CXCL12 variant prior to analyzing net BRET. Values obtained from this experiment were used to calculate the values detailed in Figure 2E and 2F.

**Supplemental Figure 3**

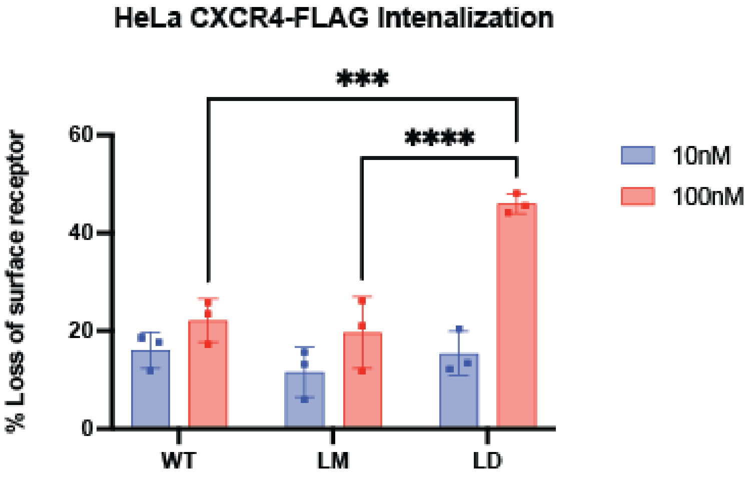


**CXCR4-FLAG transfected HeLa cells show similar internalization compared to AML cell lines.** HeLa cells transfected with CXCR4-FLAG were treated with either 10 nM or 100 nM of CXCL12-WT, CXCL12-LM, or CXCL12-LD. Surface levels of CXCR4 were detected and quantified using an anti-FLAG antibody and normalized to vehicle treated cells. Significance is by two-way ANOVA from 3 independent biological replicates.

**Supplemental Figure 4**

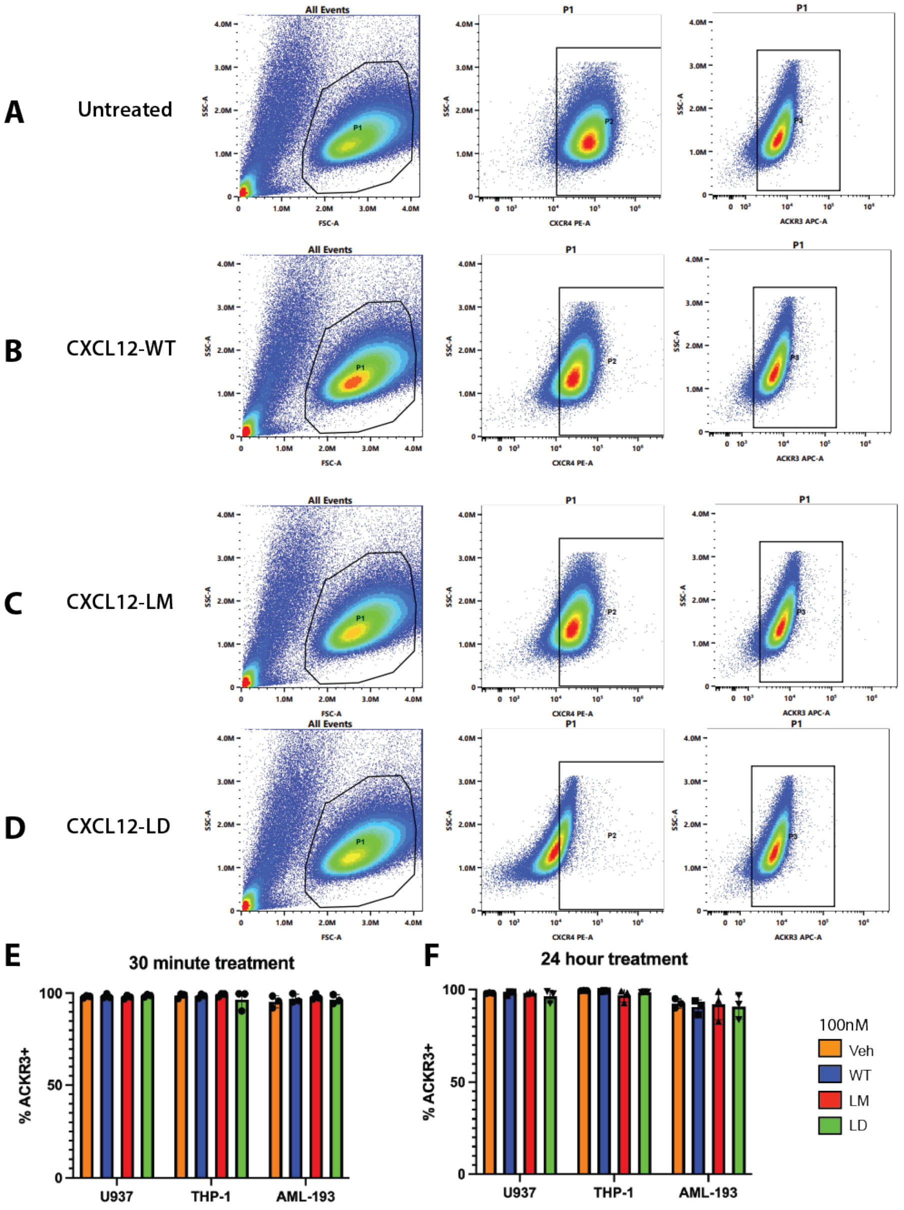


**Surface ACKR3 levels do not change with CXCL12 oligomers. (A)** Gating scheme for single cells (left column), CXCR4 surface staining (middle), and ACKR3 surface staining (right). U937 cells were untreated, **(B)** treated with CXCL12-WT, **(C)** CXCL12-LM, **(D)** or CXCL12-LD for 24 hours. Flow cytometry scatter plots representative of 3 independent biological replicates. **(E-F)** Quantified ACKR3 surface staining on U937, THP-1, and AML-193 cell lines after treatment with the indicated CXCL12 isoform for 30 minutes **(E)** or 24 hours **(F)**. No differences in the percent ACKR3 positive cells were detected by one-way ANOVA at either timepoint in each cell type.

**Supplemental Figure 5**

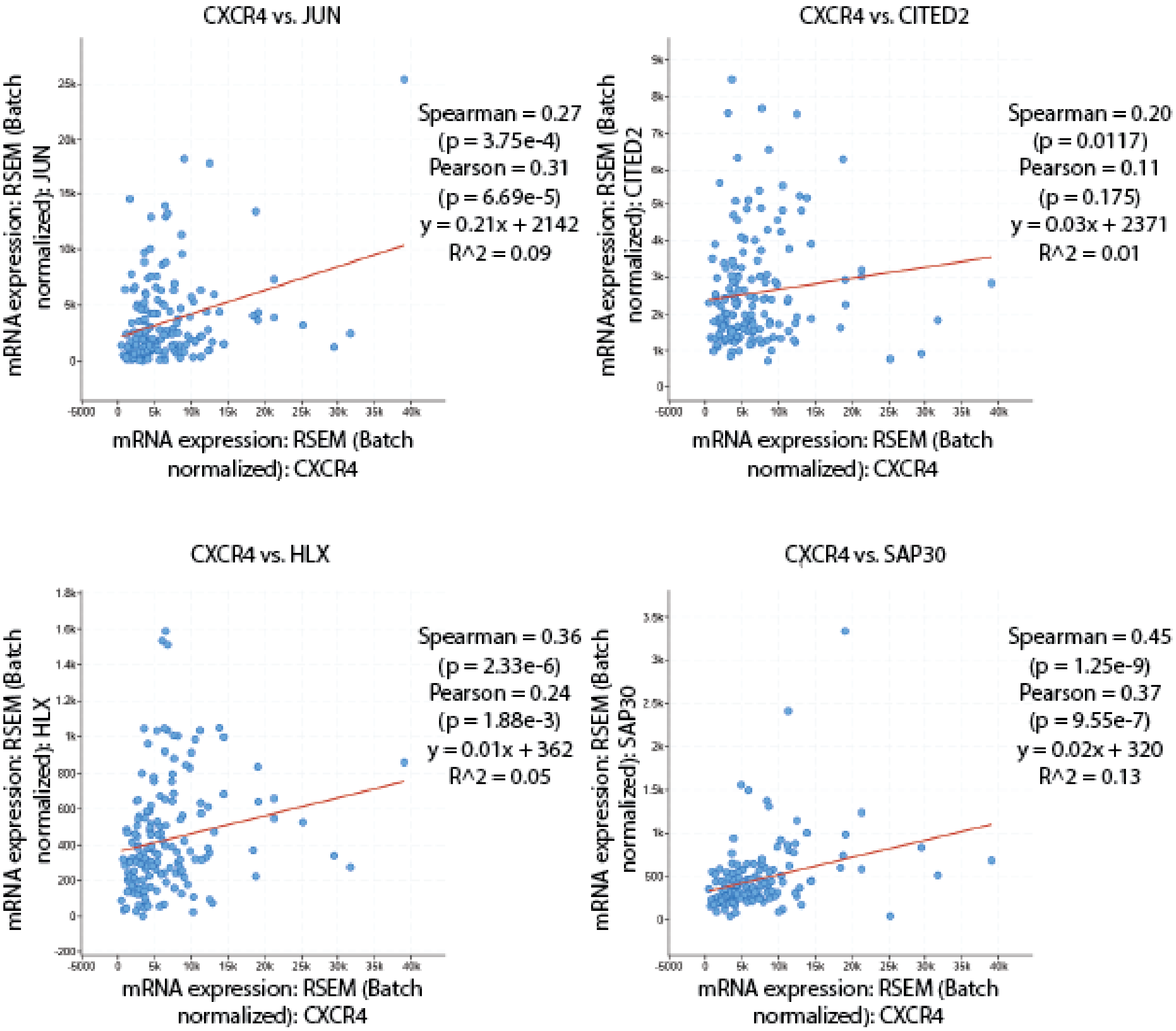


**Gene correlations with CXCR4 from The Cancer Genome Atlas Data.** cBioPortal was used to confirm the selected genes seen in Fig. 7C are correlated with CXCR4 expression in AML using The Cancer Genome Atlas AML dataset selected from the Pan-Cancer dataset. The genes include JUN (top left), CITED2 (top right), HLX (bottom left), and SAP30 (bottom right). All genes shown had a q-value, derived from the Benjamini-Hochberg FDR correction procedure, below 0.05 and a positive Spearman's correlation.

**Supplemental Figure 6**

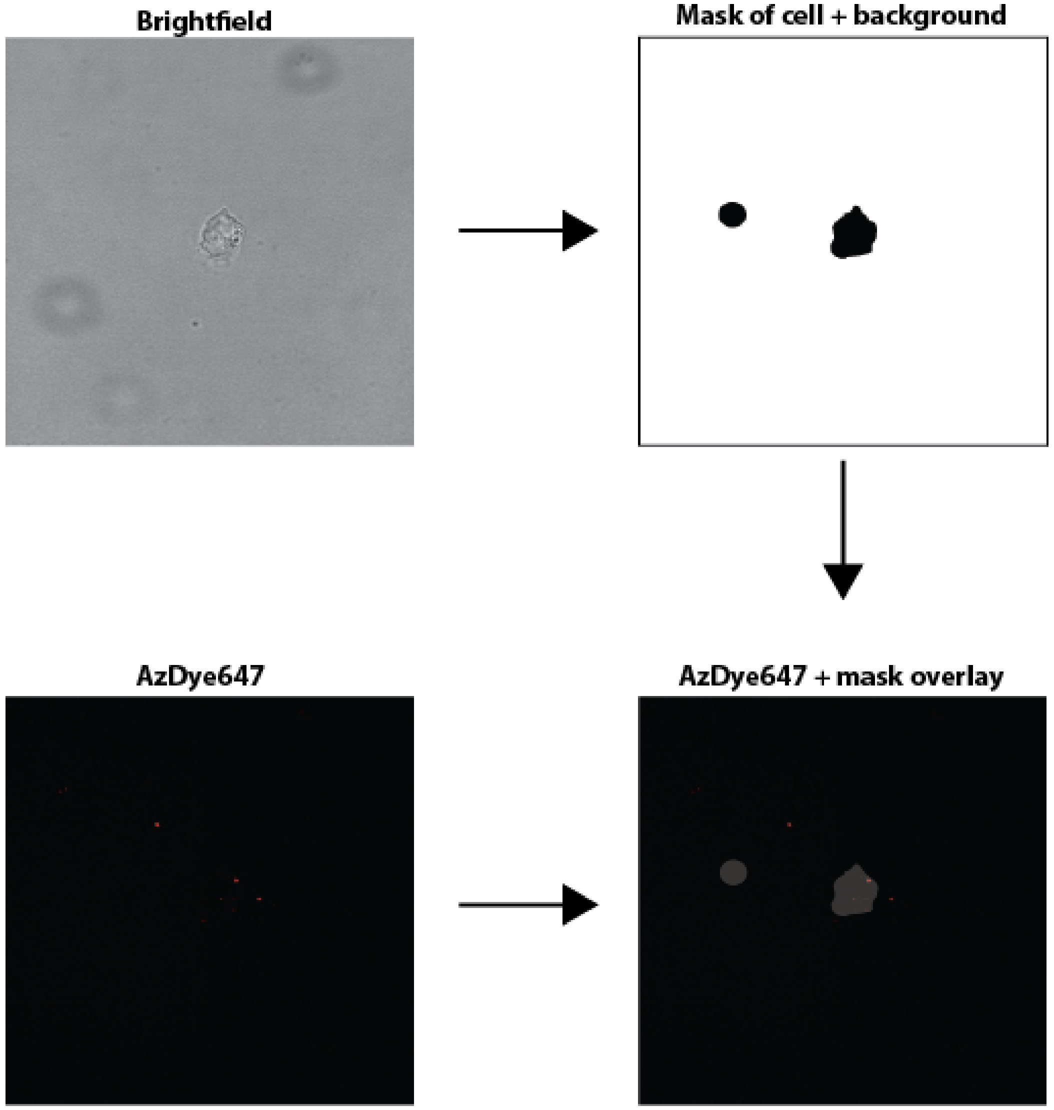


**Methodology for AzDye647-CXCL12 quantification.** The top left image is an AzDye647-CXCL12-LM treated cell, shown in brightfield at 100X. The cell trace to generate the mask of the cell is shown on the top right. The circle mask (left object within top right image) is the area taken to subtract the background staining. The mask was then overlayed onto the red-only channel image (bottom left) to generate the AzDye647 + mask overlay image (bottom right). Using this, the average AzDye647 intensity within the background mask was subtracted from the average AzDye647 intensity within the cell mask to receive the corrected average fluorescence per cell. These values were then used for the quantification shown in Figure 5D.
